# Soft, Transparent and Bioresorbable Microelectrode Array for Transient Electrophysiological Recordings

**DOI:** 10.64898/2025.12.08.692913

**Authors:** Mohamed K. M. Abdelbaki, Clément Cointe, Dina N. Arvanitis, Christian Bergaud, Ali Maziz

## Abstract

Transparent microelectrode arrays that enable multimodal investigation of spatiotemporal electrophysiological activity are critical tools for advancing the understanding of excitable tissues such as the brain, heart, and peripheral nerves. Traditional implantable devices are engineered for chronic use but require surgical removal when they fail or are no longer needed. In contrast, bioresorbable systems that naturally dissolve after serving temporary functions offer a compelling alternative, eliminating the risks and costs of extraction procedures. Here, we present the design, fabrication, and validation of a soft, fully bioresorbable, and optically transparent MEA platform for transient, bidirectional interfacing with living tissues. The device provides high-resolution electrical mapping of dynamic activity. We report precise characterization of electrochemical performance, mechanical properties, bioresorption kinetics, and biocompatibility. While validated in models of cardiac function, this platform establishes a versatile foundation for bioresorbable electrophysiological technologies with applications ranging from postsurgical monitoring of transient conditions to the study and treatment of neurological and neurodegenerative disorders.

## 1. Introduction

Implantable bioelectronic systems have become indispensable tools for advancing our understanding of excitable tissues and guiding clinical interventions across diverse organ systems, such as the brain, heart, or peripheral nerves ^[1]^. By enabling high-resolution spatiotemporal recordings and stimulation, these devices provide unique insights into physiological and pathological processes and have transformed both basic research and medical practice. With implanted microelectrode arrays (MEAs), it is possible to target well-defined anatomical regions, enabling localized recording and/or delivering electrical signals at the cell level ^[2,3]^. Implanted MEAs enable high-resolution mapping and monitoring of neural activity to guide resective neurosurgeries such as those performed for epilepsy ^[4]^ or brain tumors ^[5]^, and facilitate precise neurodevice placement in movement disorders like Parkinson’s disease ^[6]^, assist surgical interventions involving complex or highly interconnected neural structures, support the assessment of cardiac conduction abnormalities, and advance investigations of peripheral nerve function ^[7]^. While the information provided by these platforms has been invaluable, their design has historically been optimized for chronic use, leading to persistent challenges related to long-term stability, material biocompatibility, and the risks associated with device explantation once they are no longer needed ^[8–10]^. Traditional implantable electrodes are typically constructed from mechanically rigid and chemically stable materials, such as silicon, platinum, and iridium oxide. These devices have enabled precise mapping of electrophysiological activity since the pioneering work of silicon-based probes ^[11]^. However, the inherent stiffness of such systems leads to mechanical mismatch with soft and dynamic biological tissues, causing foreign body reactions, glial scarring in the brain, fibrosis in cardiac or neural tissues, and eventual signal degradation ^[12]^. Furthermore, their permanence introduces additional complications, including chronic immune responses, the risk of infection, and the necessity of invasive surgical procedures for removal ^[13]^. These factors limit their clinical applicability, particularly when temporary monitoring is required, as in post-operative evaluation or transient pathological conditions ^[14]^. Recent efforts have focused on addressing these limitations through the development of soft and flexible bioelectronic interfaces ^[15]^. Advances in materials science and microfabrication have enabled electrodes constructed from polymeric substrates such as parylene ^[16–19]^, polyimide ^[20,21]^, or hydrogels ^[22]^, which reduce mechanical mismatch and improve integration with living tissues. Similarly, transparent conductors and polymers, including graphene ^[23]^ and PEDOT:PSS ^[24,25]^, have facilitated multimodal approaches by enabling simultaneous optical imaging and electrophysiological recordings. These innovations have extended the performance and utility of implantable devices, yet they remain constrained by their permanent nature. Ultimately, permanent devices still require removal and expose patients to additional surgical risks ^[10,12,14]^.

In parallel, the emergence of bioresorbable medical devices has introduced a paradigm shift in transient bioelectronics ^[26,27]^. By leveraging biodegradable polymers such as silk fibroin, poly(lactic acid) (PLA), poly(lactic-co-glycolic acid) (PLGA), and polycaprolactone (PCL) ^[27]^, along with hydrolysable metals and semiconductors such as magnesium, zinc, molybdenum, and silicon, researchers have demonstrated novel platforms that naturally dissolve after serving a temporary function ^[28]^. These materials dissolve in the body after device implantation through various mechanisms and generate safe degradation by-products ^[29–32]^, eliminating the need for surgical extraction and thus reducing patient risk and healthcare burden ^[32]^. Bioresorbable devices have already found success in areas such as drug delivery and temporary vascular stents, and recent progress has extended their use toward transient bioelectronic interfaces for electrophysiological monitoring ^[33,34]^. Initial demonstrations have established feasibility, but many of these systems are constrained by short functional lifetimes, limited spatial resolution, or insufficient mechanical and optical properties for advanced multimodal studies ^[13,35–37]^. Despite these advances, there remains an unmet need for implantable bioelectronic systems that combine softness, optical transparency, and bioresorbability within a single platform. Soft mechanics are critical to minimize tissue damage and immune response, transparency is essential for enabling combined optical and electrical investigations of excitable tissues, and bioresorbability ensures safe disappearance after the device has fulfilled its purpose, eliminating the risks associated with permanent implantation. Integrating these features into a single system would represent a significant advance for both fundamental research and clinical practice, providing transient yet high-performance monitoring capabilities for conditions ranging from post-surgical neural surveillance to cardiac arrhythmia mapping and peripheral nerve assessment.

Here, we report the design, fabrication, and validation of a fully bioresorbable, soft, and transparent microelectrode array capable of transient electrophysiological recordings (**Figure 1**). This platform integrates biocompatible and biodegradable materials to provide high-resolution electrical mapping of dynamic activity while maintaining optical transparency for multimodal studies. We characterize its electrochemical performance, mechanical properties, bioresorption kinetics, and biocompatibility, and we validate its functionality in 3D models of cardiac physiology. By uniting softness, transparency, and bioresorbability, this work establishes a versatile foundation for next-generation transient bioelectronic systems with broad applicability across neural, cardiac, and peripheral nerve interfaces.

**Figure 1.**
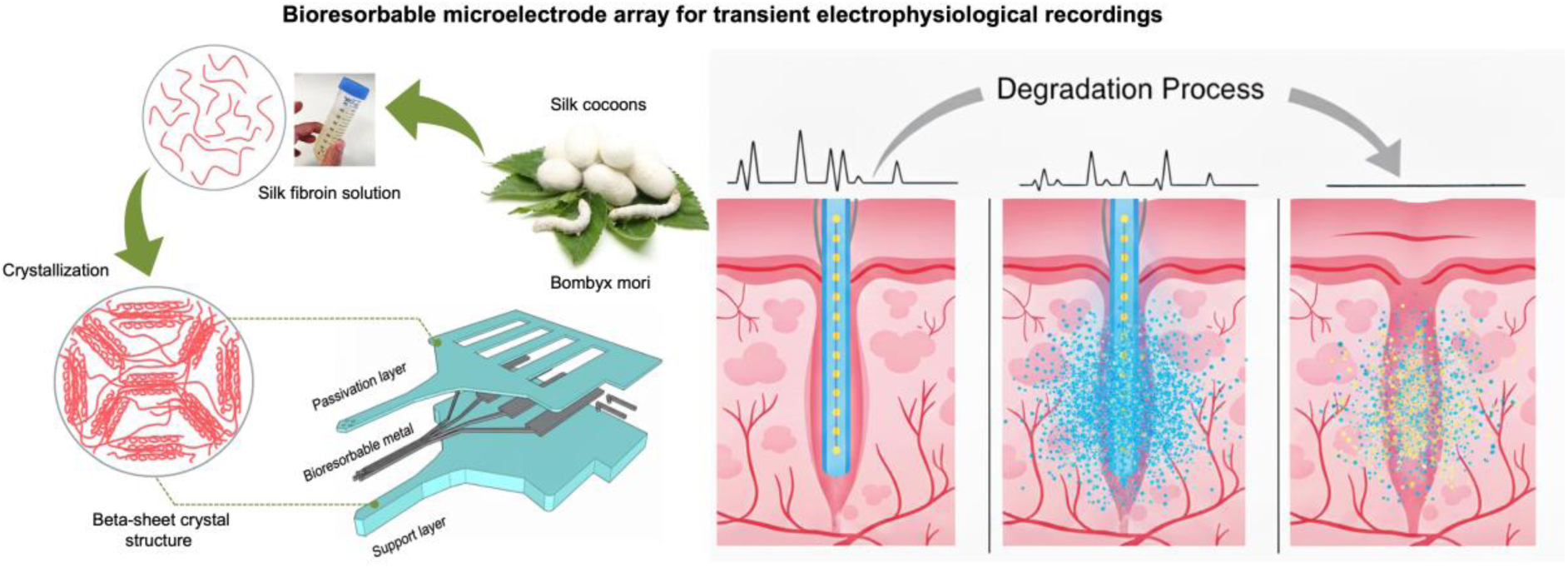
Schematic of a soft, optically transparent, and fully bioresorbable microelectrode array. Left: The fabrication process begins with fibroin protein extracted from Bombyx mori silk cocoons, which is microfabricated into a multilayer bioresorbable electrode array (substrate, metal traces, and passivation layers). Right: Illustration of the device’s life cycle, showing high-quality electrophysiological recordings at t = 0, progressive loss of function as the materials degrade, and complete dissolution at end-of-life.

## 2. Results and Discussions

### 2.1. Design and fabrication of bioresorbable and transparent microelectrode array

Our microelectrode array consists of a silk substrate base, a patterned iron (Fe) electrode array, and a silk encapsulation layer. The microelectrode array includes multiple microelectrode sites for tissue contact, interconnecting channels, and connector pads for external instrumentation (**Figure** 2A, C-E). Silk fibroin was chosen for its excellent biocompatibility, mechanical flexibility (elastic modulus **≈** 3 GPa) ^[18,38]^, and controllable degradation profile, which makes it an ideal material for the development of the proposed transient MEA platform ^[39]^. Additionally, the transparency of silk fibroin also facilitates optical imaging in parallel with electrophysiological recordings. Silk fibroin degrades into non-harmful peptides and free amino acids that can be easily removed in biological media. The lifetime of the bioresorbable silk layer can be specifically adjusted via the crystallinity of the silk protein, i.e., the β-sheet content. In particular, treating silk by immersion in polar solvents (e.g. methanol, ethanol, etc.) increases the β-sheet content in the film ^[40,41]^. For instance, in a proteolytic medium (protease XIV (PXIV) in PBS solution (1 U/ml)), silk fibroin samples degrade at different rates depending on the methanol treatment time. We found that films treated for 3 and 6 h degrade within a few hours while those treated for 12 and 24 h last up to four weeks (**Figure S1A**).

**Figure 2.**
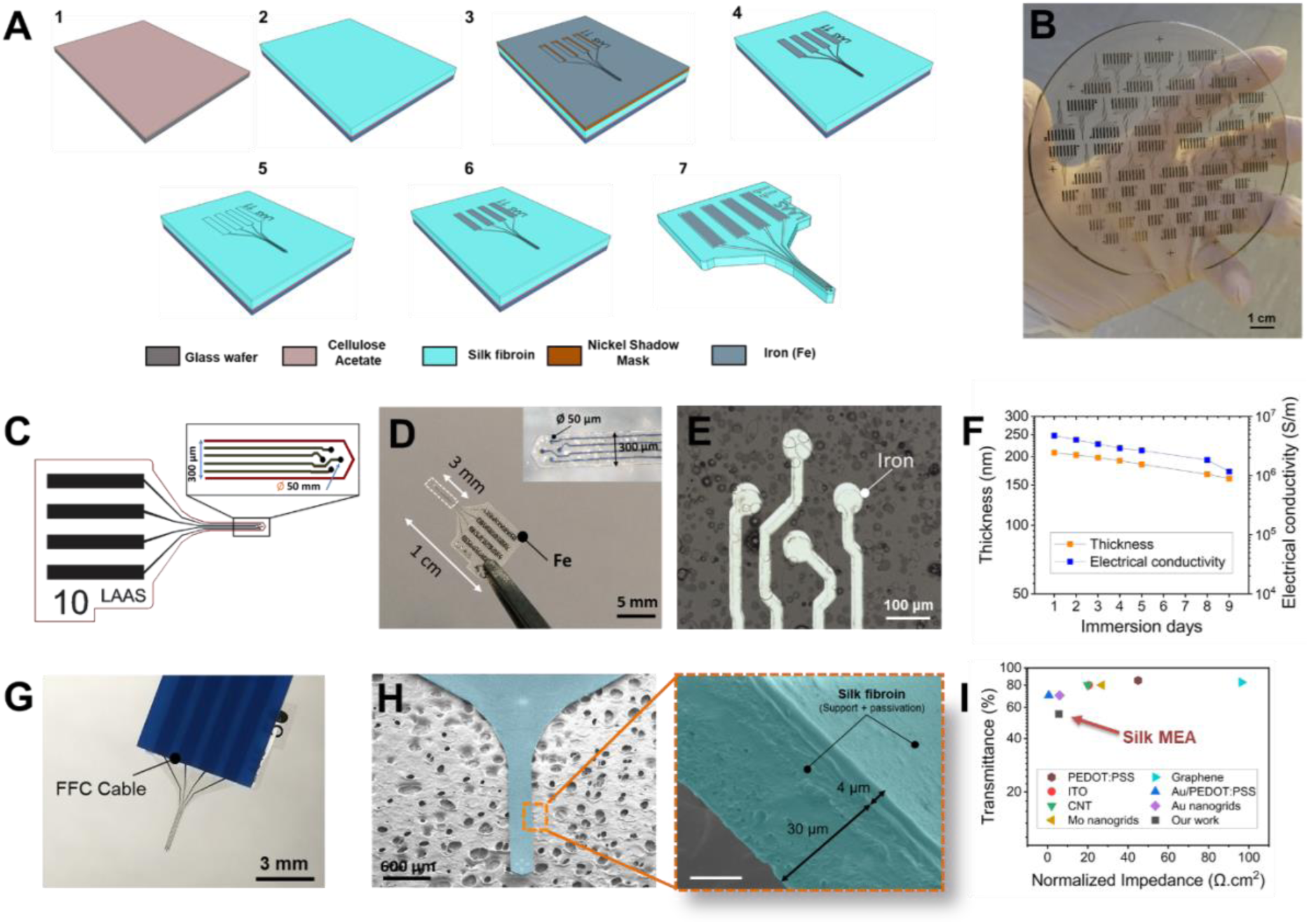
Fabrication of bioresorbable MEAs. **A**) Schematic illustration of the main fabrication steps on glass substrate: (1) CA deposition, (2) silk fibroin support deposition, (3) Fe deposition with as Ni shadow mask, (4) Patterned Fe microelectrodes, (5) silk fibroin passivation deposition, (6) Etching of electrode openings by RIE and (6) Final shaping of the silk MEA by laser etching and release from the substrate. **B**) Scalable batch microfabrication of silk MEA on a 4-inch glass wafer (54 elements per wafer). **C**) Schematic illustration of the MEA with a total footprint of 1 cm and a 3 mm-long and 300 μm-wide implantable shank of featuring four 50-µm-diameter recording microelectrodes. **D**) Picture of the microfabricated MEA, the inset shows the corresponding zoom-in on the implantable shank with 4 Fe microelectrodes. **E**) Optical microscopy image of the patterned Fe deposited by PVD. **F**) Evolution of the electrical conductivity and the corresponding thickness of Fe electrodes over degradation in ACSF solution at 37 °C for 8 days. **G**) Image of the bioresorbable MEA showing the Fe contact pads connected via an FFC cable, used for subsequent electrochemical and electrophysiological measurements. H) SEM picture of the final MEA shank, the inset highlights the strong adhesion between the support and passivation silk layers. **I**) Comparison of normalized impedance and transmittance of the fabricated silk MEA compared to various transparent MEA materials used in electrophysiological research.

The Fe electrodes were selected as the conductive material due to their bioresorbable nature, favorable corrosion resistance in physiological environments, and mechanical robustness compared to other biodegradable metals ^[42]^. This combination ensures stable signal acquisition while enabling safe resorption after the functional period. Fe gradually dissolves into non-toxic ionic products in aqueous environments according to the reaction:2Fe+ O_2_ + 2H_2_O→ 2Fe(OH)_2_. The dissolution kinetics and thickness evolution of 200-nm-thick Fe films were investigated in ACSF solution (pH 7.4, 37 °C). The measured dissolution rates were 6.07 ± 0.28 nm per day at 37 °C. At physiological temperature (37 °C), approximately 0.37 ng of Fe dissolves per day, which remains well below the tolerable daily intake level (∼10-15 mg/day) ^[43–45]^. In addition, the initial electrical conductivity of Fe was measured to be (4.74 ± 0.05) × 10⁶ S.m⁻¹ before degradation, gradually decreasing over time to (1.17 ± 0.06) × 10⁶ S.m⁻¹ at day 9 (Figure 2F). This trend reflects the progressive thinning of the metallic layer while maintaining sufficient electrical conductivity. These results confirm that Fe exhibits a controlled and biocompatible degradation profile suitable for transient bioelectronic applications.

Technological developments combine several microfabrication techniques, including silk spray-coating, soft lithography, etching, and Fe deposition through fine-line stencil masks (further details are provided in the Materials and Methods section and the Supporting Information). The final shaping of the microelectrode array is performed by reactive ion etching using a 75/25 O_2_/CF_4_ gas ratio (**Table S1**). These etching conditions enable the fabrication of silk fibroin structures with precise control over geometry, dimensions, and overall shape, as illustrated in Figure 2A.

The proof-of-concept device shown in Figure 2B integrates 54 elements, each consisting of four to eight Fe recording/stimulation microelectrodes (50μm in diameter) patterned onto probes measuring 3mm in length and 300μm in width. This demonstrates that large-scale batch fabrication of well-defined silk microelectrode arrays can be reliably achieved directly on glass substrates.

Transparency of the final device is also essential for enabling combined optical and electrical investigations of excitable tissues. The average optical transmittance of the silk-based bioresorbable MEA was measured at 55% across the visible spectrum, comparable to or exceeding previously reported values for other transparent MEA materials used in electrophysiological studies (Figure 2I) ^[46–52]^. Overall, the bioresorbable microelectrode array exhibit an excellent balance between electrical functionality, mechanical compliance, and optical transparency, making them particularly suitable for transient multimodal investigations of neural activity and tissue responses.

### 2.2. Evaluation of mechanical properties

Investigating the mechanical behavior of our bioresorbable MEA is crucial to ensure both their structural stability during implantation and their controlled, safe degradation within biological tissues. In particular, assessing their response to compressive stress offers critical insights into how the devices withstand insertion forces and maintain intimate contact with tissue without mechanical failure. We first investigated the mechanical performance of our MEA under axial compression against a rigid substrate. Slender beams ultimately undergo buckling once a critical load (*F*_buckling_) is exceeded. As shown in **Figure 3**A, compression tests conducted on a rigid substrate demonstrate the clear mechanical advantage of the silk assembly. The devices exhibited robust strength, with an average buckling force of 16.55 ± 0.59 mN, which is approximately 10-fold higher than the penetration force required for insertion into tissue-mimicking gel (Figure 3B). These experimental results were further compared with theoretical predictions (Figure 3C). The silk device can be modeled as clamped beams, where the critical buckling force is described by Euler’s formula: *F*_buckling_ = *π^2^I_x_E/(KL)*^2^ ^[53]^. Here, *E*is the Young’s modulus of the material, *I*_*x*_ = *wt*^3^/12is the area moment of inertia along the x-axis, and *L, w*, and *t*denote the length, width, and thickness of the probe, respectively. The effective length factor was taken as *K* = 0.7, consistent with beams fixed at one end. Assuming a Young’s modulus of 3 GPa for silk fibroin ^[18,54]^, the theoretical model yields a buckling force of 7.2 mN for 35 μm-thick devices, which closely aligns with both the order of magnitude and the trend observed experimentally (Figure 3A).

**Figure 3.**
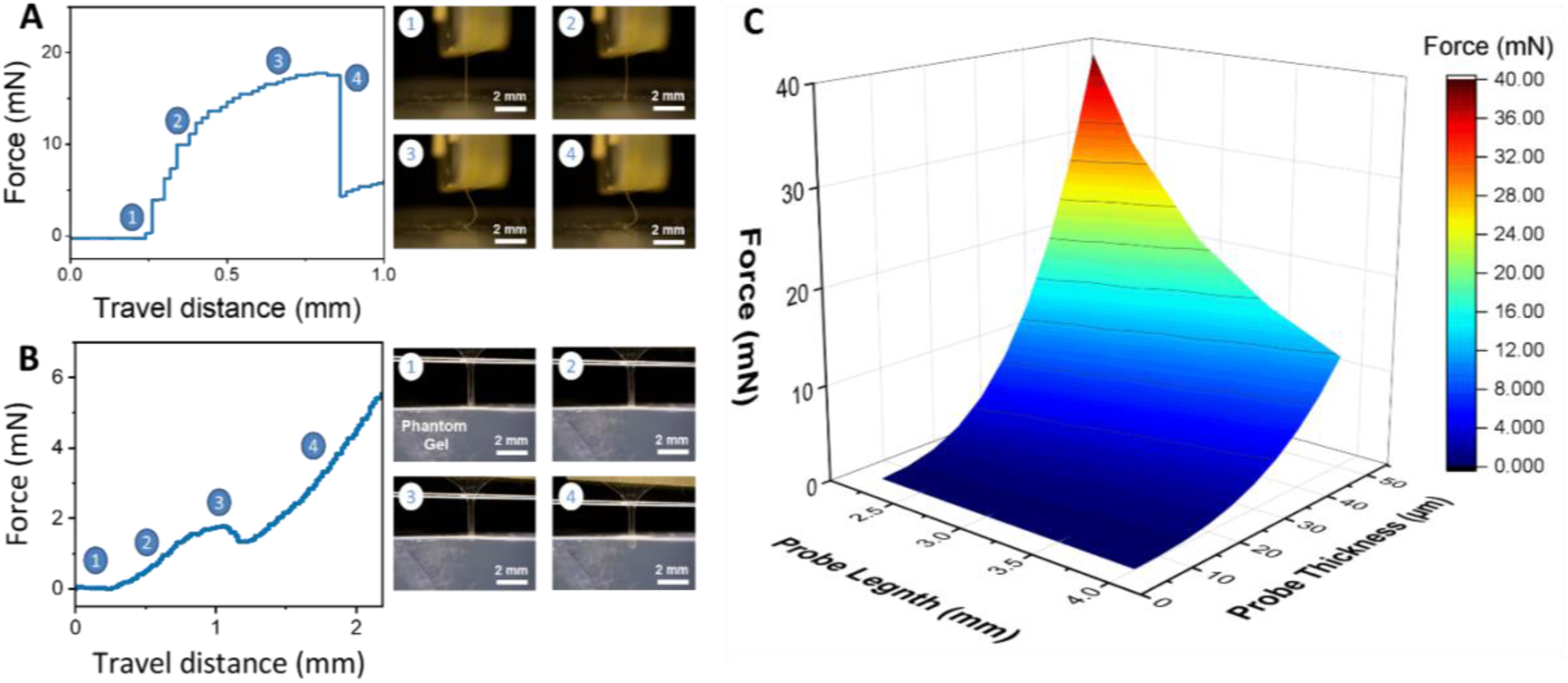
Mechanical characteristics of bioresorbable MEAs. **A**) Compression force profile of MEA probes against a hard substrate, optical pictures illustrating the corresponding stages of the shank compression in correlation with the force evolution. **A**) Compression force profile of the MEA probe against a hard substrate, optical pictures illustrating the corresponding stages of the shank compression in correlation with the force evolution. **B**) Insertion test of the MEA probe in brain phantoms. The force profile was recorded during the shank’s insertion into a 1 wt% agarose phantom gel. Optical images depict the various insertion stages relative to the force evolution. **C**) Theoretical modelling of the MEA probe showing the effect of shank length (mm) and thickness (µm) on buckling force. The model assumes a clamped–pinned configuration, with a constant rectangular cross-section. The shanks are treated as beams fixed at one end and pinned at the other (K = 0.7), where the Young’s modulus (E) of silk fibroin is 3 GPa.

We further assessed the insertion performance of the devices through cyclic implantation into a tissue-mimicking phantom composed of 1 wt.% agarose gel. Figure 3B illustrates the sequential process of probe insertion: (1) the probe mounted on the manipulator approaching the gel, (2) initial contact with the gel surface, (3) successful perforation of the gel, and (4) deep insertion into the gel. The MEA penetrated the phantom consistently without any observable buckling or bending, demonstrating that the insertion shuttle possesses sufficient stiffness for reliable implantation. From the insertion force profile (Figure 3B), the first peak force observed during perforation (stage 3) was 1.77 mN, representing the minimum force required to penetrate the tissue-mimicking gel. This value aligns well with previously reported insertion forces for similar systems ^[38]^. The tissue phantom used in this study is a standard model for mimicking electrode implantation in gray and white matter ^[55]^. However, it does not capture several important anatomical features of the brain. In practice, the minimum insertion force varies with multiple factors, including the animal species, the specific tissue layer (pia, dura mater, gray or white matter), the geometry of the shank, and the insertion speed ^[55]^. For instance, penetrating the dura mater requires significantly higher forces. Our fabrication approach offers full flexibility in material stiffness, dimensions, and shank geometry, enabling us to precisely tailor the device’s buckling resistance to the intended biological target.

### 2.3. Electrochemical bench testing of bioresorbable microelectrode array

Electrochemical impedance spectroscopy (EIS), cyclic voltammetry (CV), and degradation kinetics were performed to assess the interfacial properties and stability of the bioresorbable MEAs over time. These measurements provide insight into the charge transfer capacity, double-layer behavior, and signal fidelity of the transient electrodes under physiologically relevant conditions._**Figure 4** A, F shows the impedance spectra of the electrodes recorded from day 1 (d1) to day 9 (d9). At d1, the electrodes exhibited relatively low impedances across the frequency spectrum, with values around ∼275-300 kΩ at 1 kHz, which is suitable for electrophysiological recordings. Over time, the impedance increased progressively, reaching ∼1.06–1.23 MΩ at d9. This increase can be attributed to ongoing hydrolytic corrosion and partial dissolution of the Fe microstructures, which reduce the effective conductive area and increase the interfacial resistance. The frequency-dependent impedance drop at high frequencies (>10^3^ Hz) reflects the capacitive nature of the electrode–electrolyte interface, whereas the stabilization at low frequencies suggests charge-transfer resistance dominance. Importantly, despite the degradation, the impedance remained within the range typically considered acceptable for tissue interfaces during initial days, highlighting the potential of these devices for electrical recordings.

**Figure 4.**
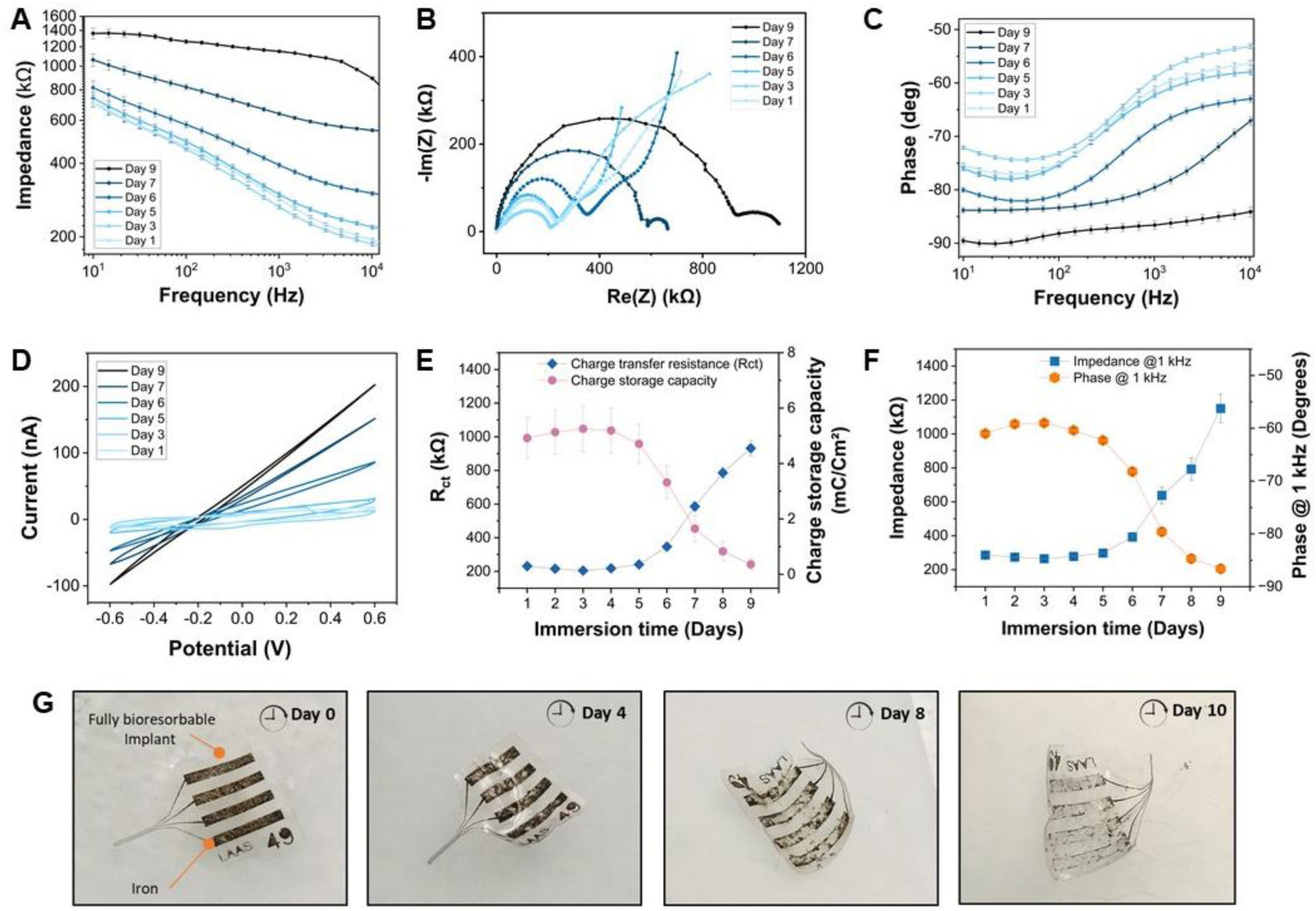
Electrochemical measurements of Fe microelectrodes as a function of degradation in PBS gels (agarose gels with PBS 0.5 wt%). **A**) EIS measurement of Fe microelectrodes over a frequency range of 10 Hz to 10 kHz. **B**) Corresponding Nyquist plots of Fe microelectrodes as a function of frequency. **C**) Phase plots of Fe microelectrodes versus frequency. **D**) Cyclic voltammetry recorded at 200 mV/s within a potential window of ±0.6 V versus Ag/AgCl reference electrode. **E**) Evolution of the charge transfer resistance (R_ct_) in relation to the charge storage capacity (CSC) during Fe microelectrode degradation. **F**) Evolution of the impedance in correlation with the phase angle of Fe microelectrodes. **G**) Enzymatic degradation of bioresorbable MEAs in PBS containing protease XIV (1 U/ml), with photographs illustrating the progressive degradation and loss of non-passivated metal tracks in fully bioresorbable electrodes.

The corresponding phase spectra (Figure 4C, F) reveal initially strong capacitive behavior, with phase angles close to –61° at 1 kHz, gradually shifting toward less negative values (∼ 86°) over time. This shift indicates a progressive loss of capacitive dominance and an increasing contribution of resistive processes as the electrodes corrode. Nyquist plots (Figure 4B) further confirm this trend: the semicircle diameters associated with charge-transfer resistance (R_ct_) increased substantially during the degradation period, reaching ∼900–980 kΩ by day 9. Extracted R_ct_ values (Figure 4E) show a steady rise after day 5, consistent with a more resistive electrode–electrolyte interface. Such behavior is expected for dissolving metallic systems and provides further evidence of a temporal window during which the electrodes preserve favorable capacitive properties for neural recording.

Cyclic voltammetry (Figure 4D) provides additional insight into the electrode’s charge storage ability. At day 1, the CV curves displayed well-defined capacitive features with anodic and cathodic currents exceeding 20 nA, consistent with efficient reversible charging and discharging of the double layer. Over time, the CV envelopes progressively narrowed, and the overall current increased, confirming the loss of effective surface area and charge injection capacity. Quantitative analysis of the charge storage capacity (CSC, Figure 4E) showed stable values during the first 5 days, followed by a pronounced decline after day 6, in agreement with the impedance and R_ct_ evolution. This reduction in CSC is a direct consequence of the electrode degradation process, ultimately leading to the loss of electrochemical activity. Nevertheless, the charge transfer capabilities of the bioresorbable microelectrodes remained compatible with neural signal detection over the relevant time frame of several days.

### 2.4. Cytotoxicity assessment

To assess the cytotoxicity of the devices, cell viability assays were performed demonstrating that the fabricated microelectrode array maintains high cellular viability throughout the degradation process. As shown in **Figure 5**, SH-SY5Y neuroblastoma cells were incubated with the devices for 24 h, 1 week, 2 weeks, and 3 weeks at 37 °C. MTT assay results revealed no significant decrease in metabolic activity at any time point, confirming the absence of cytotoxic effects. These findings indicate the excellent non-toxicity of both the silk fibroin substrate-encapsulation layers, and the Fe-based conductive components. Moreover, the degradation by-products generated over the incubation period did not induce any measurable toxic response, suggesting that the silk-Fe system provides a safe transient platform suitable for long-term biological interfacing.

**Figure 5.**
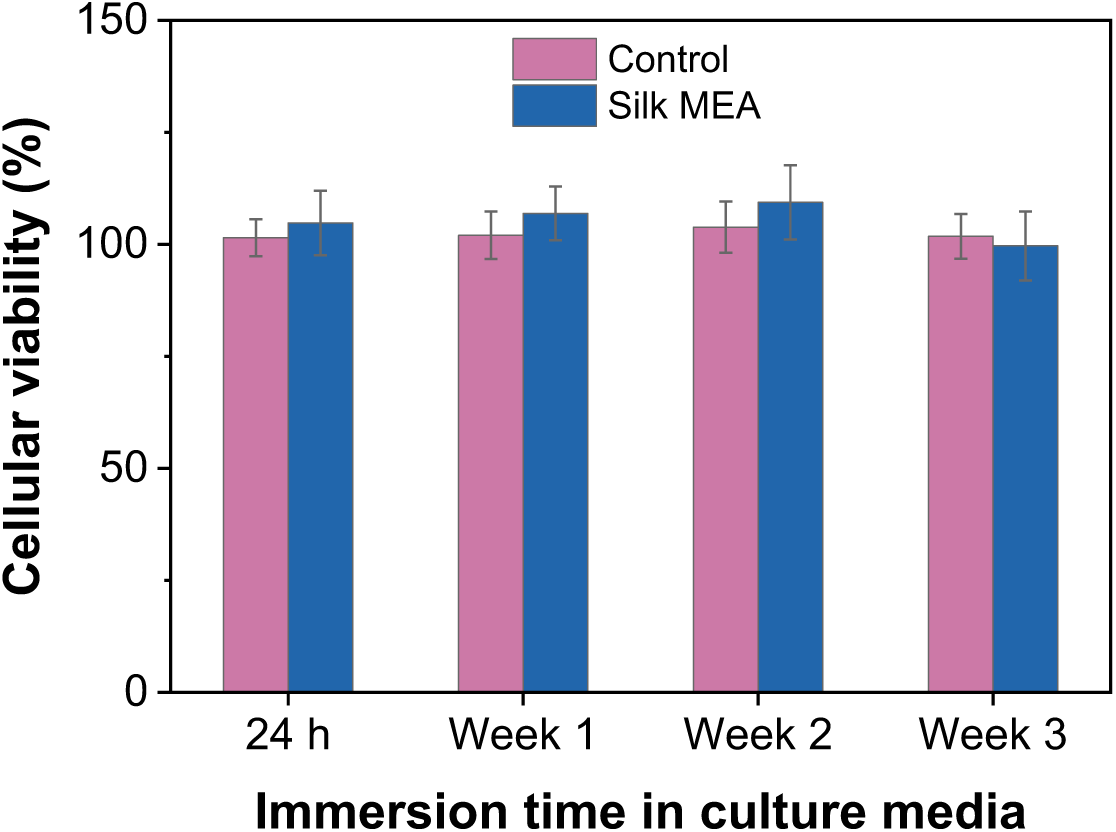
Cytotoxicity assessment of bioresorbable MEAs. MTT elution test results for fully bioresorbable silk MEA and coverslips with biocompatible glue only as a control. Cell viability is expressed relative to control solutions.

### 2.5. Ex vivo demonstration in cardiac 3D organoid models

In vitro experiments on human pluripotent stem cell (hPSC)-derived complex 3D cardiac organoids demonstrate the functionality and capability of the developed bioresorbable MEA for spontaneous, localized electrical interrogation of cardiac electromechanical function. **Figure 6**A illustrates the experimental setup for interfacing with 3D cardiac organoids, where the microelectrode array is positioned underneath the 3D organoid, allowing the tissue to attach directly on top of the microelectrodes, as depicted in Figure 6B. This configuration ensures intimate contact between the organoid and the recording sites, enabling stable and high-fidelity electrophysiological recordings ^[56]^. The electrical properties of 3D cardiac organoids were assessed by measuring their extracellular field potentials (FPs) over a long-term period (2 weeks). During the early recording stages, two distinct types of electrophysiological signals were detected: small, regular FPs reflecting local cardiomyocyte activity, and occasional high-amplitude bursts appearing in an apparently stochastic manner (Figure 6C, E) and Movie S1 (Supporting Information). These bursts are unlikely to originate from intracellular coupling, as the electrochemical nano-structuring of the microelectrodes surface does not enable cell penetration. Instead, they can be interpreted as transient episodes of enhanced electrode–tissue coupling or network-wide synchronization. A similar behavior was described by Yin et al. ^[57]^, who reported that the electro-mechanical signal quality of human cardiac organoids depends critically on a “sweet-spot” of mechanical contact between the organoid and the MEA, slight shifts in contact pressure produced abrupt amplitude changes and intermittent high-gain events. In our configuration, contractions of the organoid or local swelling of the soft substrate may temporarily increase the sealing resistance, which represents the electrical resistance between the cell membrane and the electrode surface and corresponds to the current leakage path. A high sealing resistance indicates a tight seal between the cell and the electrode and improves the capacitive coupling, therefore leading to the observed bursts. Their random occurrence thus reflects the non-stationary mechanical interaction between the organoid and the biodegradable electrode surface, combined with the intrinsic variability in synchronization among pacemaker clusters within the tissue.

**Figure 6.**
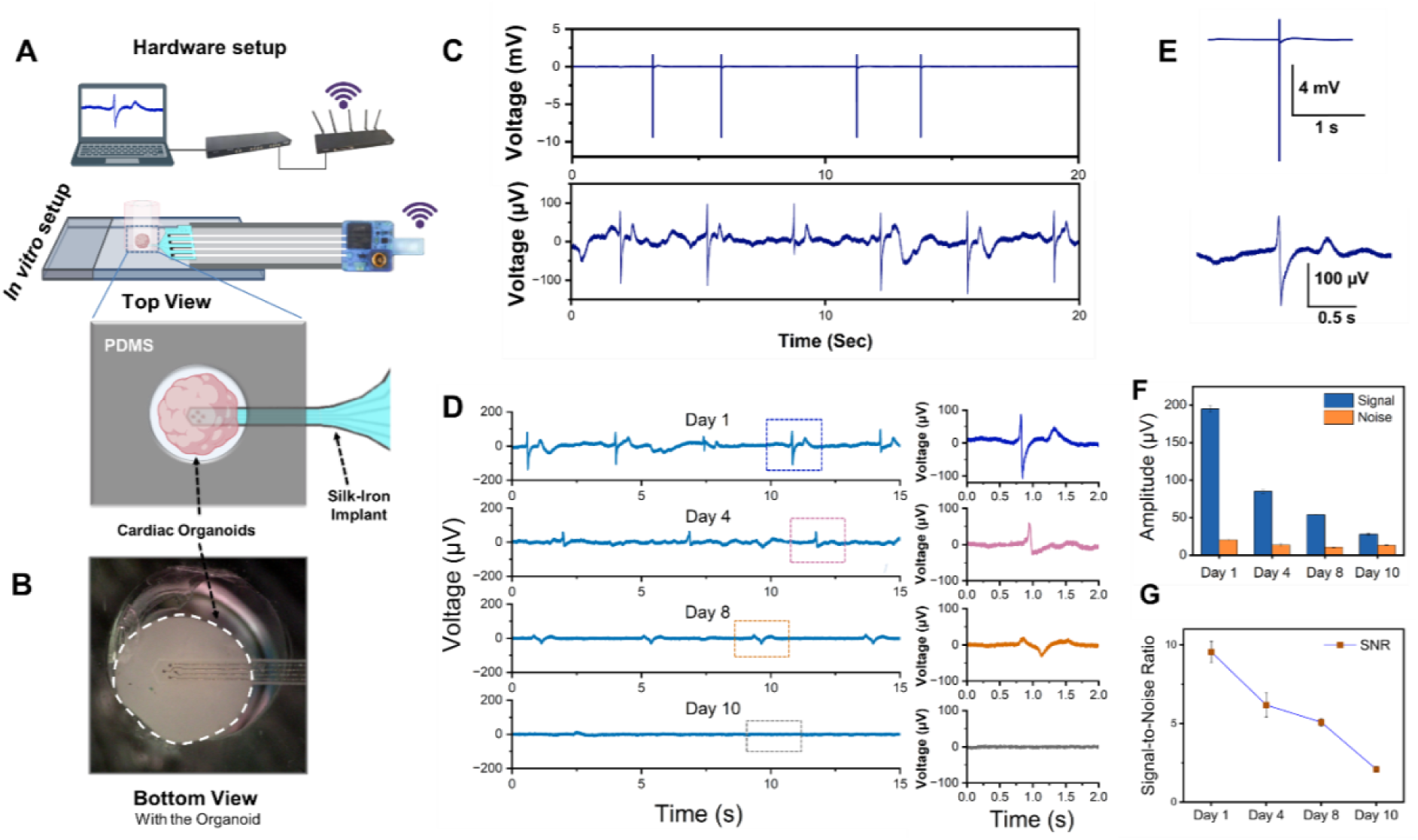
Electrophysiological recordings from 3D cardiac organoids using bioresorbable MEAs. **A)** Schematic illustration of the *in vitro* recording setup, the MEA was assembled horizontally with a PDMS mold on top containing the 3D cardiac organoid. The assembly was connected to a wireless MCS (W200-HS4) recorder. **B)** Optical image of the recording setup representing the confinement of 3D cardiac organoid on top of silk MEA. **C, E)** *In vitro* electrophysiological recordings from a hPSC-derived 3D cardiac organoid showing two types of signals: small, regular FPs (top), and occasional high-amplitude bursts (bottom). **D)** Long-term monitoring of low intensity, regular LFPs over 9 days; the initial peak-to-peak amplitude of ∼200 µV gradually decreases over days with the biodegradation of Fe microelectrodes. **F, G**) Evolution of peak amplitudes and the corresponding SNR of the recorded LFPs over days.

We further tracked the electrophysiology of a spontaneously beating cardiac organoid over two weeks and observed region-specific changes from the electrodes among recordings collected on days 1, 4, 8, and 10 (Figure 6D). Over time, as the electrode undergoes biodegradation (Silk Fibroin + Fe), its electrochemically active area and double-layer capacitance decrease, while the interfacial impedance rises. This evolution progressively reduces the efficiency of capacitive charge transfer. Consequently, only the low-amplitude extracellular potentials remain detectable, whereas the transient high-amplitude bursts disappear as the system transitions from a strongly to a weakly coupled regime. These findings emphasize that the stochastic bursts observed at early stages are signatures of temporary optimal coupling between the dynamic, contractile organoid and the soft, degradable electrode interface (**Movie S1**, Supporting Information). Additionally, we observed a consistent decrease in signal amplitude and signal/noise ratio across days 4, 8, and 10, which suggests deteriorated MEA electrical functionality, likely reflecting diminished coupling among cardiomyocytes. As the device gradually dissolves in PBS, the amplitude of the recorded FP signals becomes smaller (Figure 6F, G). Gradually, the calculated SNR decreases from 195.1 ± 4.06 µV to 85.06 ± 2.29 µV, 53.78 ± 0.63 µV, and 27.952 ± 1.317 µV after 1, 4, 8, and 10 days, respectively in PBS. Microelectrodes exhibited amplitude decreases though, potentially due to the combined effects of technical and biological factors such as increased resistance between the cells and electrodes, degradation of the Fe coating, degradation of the silk support layer localized cell stress, or uneven nutrient diffusion. Nevertheless, the stable contact between the microelectrodes and cells at specific regions of the organoid enables long-term monitoring of electrophysiological changes. This feature makes the developed silk MEAs particularly valuable for studying electrical pathways in spatially organized, heterogeneous organoids and may offer deeper insights into cardiac development, arrhythmogenesis, and cardiovascular disease pathophysiology.

## 3. Conclusion

In this study, we have demonstrated a fully bioresorbable, soft, and transparent microelectrode array capable of high-resolution, multimodal electrophysiological recordings in excitable tissues. By integrating mechanically compliant, optically transparent, and biodegradable materials, the platform addresses key limitations of traditional permanent implants, including mechanical mismatch, chronic immune responses, and the need for surgical removal. Characterizations of electrochemical performance, mechanical properties, bioresorption kinetics, and cytocompatibility confirm the device’s suitability for transient applications, while validation in 3D cardiac models highlights its capacity for accurate and reliable functional monitoring. Overall, our results establish a versatile foundation for transient bioelectronic systems, offering promising opportunities for postoperative monitoring, temporary therapeutic interventions, and multimodal studies in neural, cardiac, and peripheral nerve tissues. As a perspective, future studies will focus on implementing these devices in more complex, in vivo configurations, thereby paving the way for translational applications and further advancing the use of bioresorbable interfaces in both research and clinical settings.

## 4. Materials and methods

### 4.1. Materials

Silkworm (*Bombyx Mori*) cocoons were purchased from ECO-HOBBY LTD (Bulgaria). Dialysis membrane (MWCO 3.5 KD, Spectra/Por^TM^) was purchased from Fisher Scientific. Acetone, ethanol, sodium carbonate, and lithium bromide were purchased from Sigma Aldrich. Phosphate-buffered saline (PBS, 10 X) was purchased from Fisher Scientific. Platinum (Pt) counter electrodes and silver/silver chloride (Ag/AgCl) reference electrodes were purchased from World Precision Instruments (WPI). Human iPSC-derived cardiac organoids and media were purchased from Acrobiosystems (Switzerland). Deionized water (18 MΩ at 25 °C) was used to prepare all the solutions.

### 4.2. Extraction of bioresorbable silk fibroin aqueous solution

Silk fibroin solution was prepared from natural silkworm cocoons following a previously published protocol ^[41]^. Briefly, the cocoons were cut into small pieces and boiled in a 0.02 M sodium carbonate solution for 30 minutes to obtain degummed silk fibers. The fibers were then dissolved in a 9.3 M LiBr solution at 60 °C for 4 hours. The resulting solution was dialyzed against deionized (DI) water for 48 hours, with several water changes, to remove residual salts. Finally, the dialyzed solution was centrifuged to remove impurities, yielding the regenerated silk fibroin aqueous solution (6 wt%). The whole extraction process is illustrated in **Figure S2**.

### 4.3. Microelectrode array fabrication

The silk microelectrode arrays were fabricated using microfabrication facilities inside a cleanroom. Glass wafers (4-inch, EagleXG) were used throughout the fabrication process. The glass wafer was cleaned with oxygen plasma (800 W, 5 min) before processing. Acting as a sacrificial layer, a cellulose acetate layer (∼500 nm) was deposited by spin-coating at 2000 rpm for 30 s (5 wt% in acetone). The previously prepared silk fibroin aqueous solution (diluted to 3.5 wt%) was deposited by spray coating at a capillary flow rate of 2 ml/min, with annealing at 50 °C during deposition to ensure rapid drying. This process was repeated for six cycles, resulting in a silk support layer of ∼30.2 ± 1.3 µm thickness (**Figure S3** A,B). The thickness of the deposited silk fibroin layer was controlled by adjusting the capillary flow rate during spray coating. A 100 nm Fe layer was then deposited by thermal evaporation (ApSy-E100 thermal evaporation system) using an electroplated patterned nickel-based shadow mask (Figure S3C,D; Figure S1B). Another layer of silk fibroin (passivation layer, 5 µm) was deposited using the same spray coating technique, then transferred onto the metal layer by water annealing (-67 cmHg, room temperature, 2 h) (Figure S3E). The silk fibroin covering the device contact and electrode areas was etched using reactive ion etching with O_2_:CF_4_ plasma (75:25) at 500 W and 20 mT, aided by a silicon shadow mask (Figure S3F,G; Table. S1). Finally, the implant outlines were shaped using a UV YAG laser (355 nm, Ø 25 μm), followed by immersion in an acetone bath to dissolve the cellulose acetate sacrificial layer and release the fully bioresorbable silk-Fe devices (Figure S3H).

After fabrication, the silk MEAs were crystallized via methanol treatment to render them water-insoluble by increasing the crystalline domains in the silk fibroin, thereby allowing programmable degradation after implantation ^[18]^. The silk MEAs were placed between two metal grids to prevent shank deformation during crystallization and then immersed in methanol at room temperature for 48 hours to increase the β-sheet content of the secondary structure. Finally, the MEAs were vacuum-dried at 40 °C to remove all traces of methanol.

### 4.4. Electrochemical characterization

Electrochemical impedance spectroscopy (EIS) and cyclic voltammetry (CV) measurements were performed in saline brain phantoms (agarose gels with 0.5 wt% PBS) maintained at 37 °C over a period of 9 days. A three-electrode configuration was used, with a Pt wire as the counter electrode, a Ag/AgCl wire as the reference electrode, and the Fe microelectrodes in the MEAs as the working electrodes. All electrochemical measurements were carried out using a Bio-Logic VMP3 potentiostat inside a Faraday cage. EIS measurements were conducted by applying a 10 mV AC signal over a frequency range of 10 Hz to 7 MHz. CV was performed by sweeping the potential between −0.6 V and +0.6 V at 0.2 mV/s versus the Ag/AgCl reference electrode, enabling the evaluation of the cathodal charge storage capacity (CSCc) of the Fe bioresorbable microelectrodes during degradation.

### 4.5. Mechanical characterization

Standard compression tests were performed against a hard substrate to assess the buckling forces (the maximum force a sample can withstand before bending) of the silk MEAs under axial compression. The device was positioned between two glass slides and fixed to a MARK-10 ESM303 test bench equipped with a MARK-10 M5-012 force sensor. Compression was applied over a shank length of approximately 3 mm. Custom-developed software enabled the equipment to monitor the force in relation to the displacement of the device toward the hard substrate. Tests were performed at a speed of 2 mm·min⁻¹ to allow optimal observation of the buckling force. Real-time images of the compression test were captured using a Dino-Lite Edge camera (AF7915MZT, Dino-Lite).

In vitro mechanical insertion tests of the silk MEAs were conducted in 1 wt% agarose brain phantom gels, which mimic the mechanical properties of brain tissue. The devices were positioned and fixed in the same manner as in the compression tests. Insertion was performed at a low speed (0.5 mm·min⁻¹) to minimize damage and to optimally monitor the forces involved. Real-time images were also captured using the Dino-Lite Edge camera to correlate the different stages of insertion into the brain phantom gels with the evolution of the force.

### 4.6. Biodegradation and cytocompatibility tests

The biodegradation of the silk MEAs was evaluated using an enzymatic degradation test over 10 days. The devices were incubated at 37 °C in 1 mL of protease solution (Protease XIV from S. griseus, 3.5 U/mg, Sigma-Aldrich) in PBS (1 U/mL) as the degradation medium. Degradation was monitored daily using digital photographs. The enzyme solution was frequently refreshed to maintain enzymatic activity.

Cytotoxicity testing of the silk MEA was carried out by the elution method in accordance with ISO 10993-5. Implant samples were sterilized in 70% ethanol at 70 °C for 30 minutes, air dried, rinsed with PBS and then incubated in culture medium (DMEM supplemented with 10% fetal bovine serum, GlutaMAX, and pyruvate) at 37 °C under 5% CO₂ for 24 h, 1 week, 2 weeks, or 3 weeks to generate conditioned media containing potential degradation products. Twenty-four hours before the end of each extraction period, SH-SY5Y cells were seeded at a density of 25,000 cells/cm² into 22 mm diameter wells and maintained in media (DMEM + 10% FBS + GlutaMAX + pyruvate) for 24 h at 37 °C and 5% CO₂. At the end of the incubation, the culture medium was replaced with the conditioned medium obtained from each extraction time point, and cells were incubated for 24 h. Cell viability was evaluated using the MTT assay. Briefly, MTT reagent was added to each well and incubated for 2 h at 37 °C and absorbance was read at 570 nm using a multimode microplate reader (Varioskan LUX, Thermo Fisher Scientific).

### 4.7. Device Packaging

Prior to recording, the silk MEAs were electrically connected to a customized flexible flat cable (FFC, PP001497, Pro-power) using a conductive silver-based epoxy glue (8331D-14G, MG Chemicals), establishing an electrical connection between the FFC contacts and the device’s contact pads (Fig. 1G). Additionally, the entire device connection area was passivated with a UV-curable glue (UV625, Permabond) to protect the device from potential liquid leakage or short circuits.

### 4.8 In vitro electrophysiological monitoring

Electrophysiological activity from the silk MEA was recorded using a wireless W2100-HS4 electrophysiology system connected to the W2100 data acquisition system (Multichannel Systems, Germany) at a sampling rate of 25 kHz. The recorded signals were band-pass filtered to isolate local field potentials using a Butterworth filter from 0.1 Hz to 300 Hz, and a separate Butterworth filter from 200 Hz to 3000 Hz was applied to isolate high-frequency action potentials. Devices bearing the microelectrodes were fixed onto a glass slide with the microelectrodes facing upward. A polydimethylsiloxane (PDMS) mold with a central aperture (3 mm diameter) was placed on top of the silk shank containing four microelectrodes to confine the tissue (**Figure 6**A). Cardiac organoids, maintained in complete cardiac maintenance medium, were gently transferred into the mold and onto the electrode area. The organoids were allowed to settle for 10 minutes before recording to ensure close contact with the electrodes and stabilize their position. Baseline recordings (without organoids) were first collected to assess noise and drift characteristics. Signal preprocessing and subsequent analyses were performed in RStudio. From the filtered traces, the evolution of peak amplitude, as well as other metrics such as beat rate, signal-to-noise ratio, and field potential duration, was extracted over time in successive time windows to characterize signal stability, amplitude drift, and responses to experimental interventions.

## Author Contributions

M.K.M.A and C.C served as co–first authors and contributed equally to this work. A.M designed the study and drafted the manuscript. All the experiments were completed by C.C, M.K.M.A, A.M and D.N.A, and A.M., conceived of the study, supervised the study, interpreted the results, and revised the manuscript. D.N.A carried out biological experiments and collected data, and C.B, D.N.A and A.M interpreted the corresponding results. C.B provided statistical methodology support. All authors contributed to the critical reading of and commented on the manuscript, helped to interpret the data, and approved the final manuscript.

## Acknowledgement

Authors thank Emilie Dumarquez (LAAS, Toulouse, France) for her help in the work on 3D cardiac organoids.

## Data Availability Statement

The data that support the findings of this study are available from the corresponding author upon reasonable request.

